# High-coverage, long-read sequencing of Han Chinese trio reference samples

**DOI:** 10.1101/562611

**Authors:** Ying-Chih Wang, Nathan D Olson, Gintaras Deikus, Hardik Shah, Aaron M Wenger, Jonathan Trow, Chunlin Xiao, Stephen Sherry, Marc L. Salit, Justin M Zook, Melissa Smith, Robert Sebra

## Abstract

Single-molecule long-read sequencing datasets were generated for a son-father-mother trio of Han Chinese descent that is part of the Genome In a Bottle (GIAB) consortium portfolio. The dataset was generated using the Pacific Biosciences Sequel System. The son and each parent were sequenced to an average coverage of 60 and 30, respectively, with N50 subread lengths between 16 and 18 kb. Raw reads and reads aligned to both the GRCh37 and GRCh38 are available at the NCBI GIAB ftp site (ftp://ftp-trace.ncbi.nlm.nih.gov/giab/ftp/data/ChineseTrio/) and the raw read data is archived in NCBI SRA (SRX4739017, SRX4739121, and SRX4739122). This dataset is available for anyone to develop and evaluate long-read bioinformatics methods.

## Background

Genome In a Bottle (GIAB) is a consortium hosted by the National Institute of Standards and Technology (NIST), primarily dedicated to the development and characterization of human genomic reference materials. The consortium includes representatives from government, industry, and academia. Currently, the GIAB portfolio includes seven genomes: the pilot genome NA12878 and two son-father-mother trios (one trio of Ashkenazi Jewish descent and the other of Han Chinese descent) (Zook et al. 2016). The trio samples were selected from the Personal Genome Project with the aim of increasing reference sample diversity.^1^ The GIAB genomes have been extensively sequenced on a number of different platforms (Zook et al. 2016). The datasets have been used to generate benchmark variant calls sets for benchmarking and validating small variant calling methods (Zook et al. 2018; Krusche et al. 2018). The benchmark calls are based primarily on short-read data and cover approximately 90% of the human reference genome (Zook et al. 2016). A number of medically relevant genes are difficult to characterize using short-read sequencing data (Mandelker et al. 2016). Therefore expanding the benchmark to more challenging variants and regions using long-read sequencing technologies is of interest to the consortium and its stakeholders, including technology and bioinformatics developers, clinical laboratories, and regulatory agencies.

In an effort to expand the benchmark to more challenging variants and regions, a high-coverage long-read sequence dataset was generated for the Han Chinese Trio using the PacBio Sequel System (Pacific Biosciences, Menlo Park CA, USA). The Sequel System utilizes single molecule, real-time (SMRT) sequencing with fluorescently-labeled nucleotides. In addition to being used to help expand the benchmark to more challenging variants and regions, the dataset will be used to improve phasing of variants and produce genome assemblies. This dataset can also be used by anyone to develop and evaluate long-read bioinformatics methods.

For the GIAB Han Chinese PacBio Sequel dataset the son was sequenced to 60X coverage and parents to 30X coverage with a subread N50 of 16 - 18 kb. The raw reads and reads aligned to both the GRCh37 and GRCh38 are available at the NCBI GIAB ftp site^2^ and the raw read data is archived in NCBI SRA.

## Methods

### Experimental design

The Han Chinese GIAB trio (Table 1) samples were sequenced on the PacBio Sequel sequencing platform. The genomic DNA was used to prepare 14 sequencing libraries, 6 for the son and 4 for both the mother and father. 79 Sequel SMRT Cells were used to generate the dataset, with 46 SMRT Cells for the son, 17 for the father, and 16 for the mother.

**Table 1:** Sample names and identification numbers for GIAB Han Chinese trio. PGP ID - personal genome project identifier.

| Sample | Coriell cell line ID | NIST ID | NIST RM # | NCBI BioSample | PGP ID |
| --- | --- | --- | --- | --- | --- |
| Chinese Son | GM24631 | HG005 | RM8393 | SAMN03283350 | hu91BD69 |
| Chinese Father | GM24694 | HG006 | N/A† | SAMN03283348 | huCA017E |
| Chinese Mother | GM24695 | HG007 | N/A† | SAMN03283349 | hu38168C |

† NIST Reference Materials are not planned for the Chinese parents, but cells and DNA are available from Coriell.

### Sample Preparation

NIST RM8393 was used for HG005 sequencing libraries, and genomic DNA for HG006 and HG007 was obtained from Coriell (NA24694 and NA24695, respectively). Genomic DNA concentration was measured using the Qubit fluorimetry system with the High Sensitivity kit for detection of double-stranded DNA (Thermo Fisher, Part #Q32854), and fragment size distribution was assessed using the 2100 Bioanalyzer with the 12000 DNA kit (Agilent, Part 5067-1508). 20 µg high molecular weight genomic DNA was sheared using the Megaruptor instrument (Diagenode, Liege, Belgium) to 40kb and this was used as input into the SMRTbell library preparation. SMRTbell libraries were prepared using the Pacific Biosciences Template Preparation Kit 1.0 - SPv3 (Pacific Biosciences, Part # 101-357-000). Once libraries were completed, they were size selected using the Blue Pippin instrument (Sage Science, Beverly MA, USA) from 20kb-50kb to enrich for the longest insert lengths possible. The polymerase v2.0 binding kit (Part #101-862-200) was used to bind polymerase to SMRTbell templates. Binding complex was cleaned using the Column Clean-up kit (Pacific Biosciences, Part #100-184-100) columns before loading to remove excess polymerase and enhance loading efficiency.

### Pacific Biosciences Sequel System Sequencing

SMRTbell libraries were sequenced on the Pacific Biosciences Sequel System using version 2 SMRT Cells (Part # 101-008-000) with 10-hour movies and diffusion loading at 6-7pM on plate. Sequel Sequencing Kit 2.0 (Part # 101-053-000) - used with 39/42 SMRT Cells for son; 12/33 SMRT Cells for parental gDNA. Sequel Sequencing Kit 2.1 (Part # 101-328-600) - used in 3/42 SMRT Cells for son and 21/33 SMRT Cells for parental gDNA. Individual SMRT cell information including instrument used, date run, cell name, and cell lot is provided as Supplemental Tables 1 - 3.

### Sequence Data Processing

Sequence data was exported from SMRT Link (version 5.0.1.9585) as tar.gz files using the “Export Data Sets” functionality. Each movie has one tar.gz file that contains sequence data in subreads BAM format and metadata (Figure 1). FASTA files were extracted from subread BAMs using samtools (version 1.3.1, Li et al. 2009)

**Figure 1:**
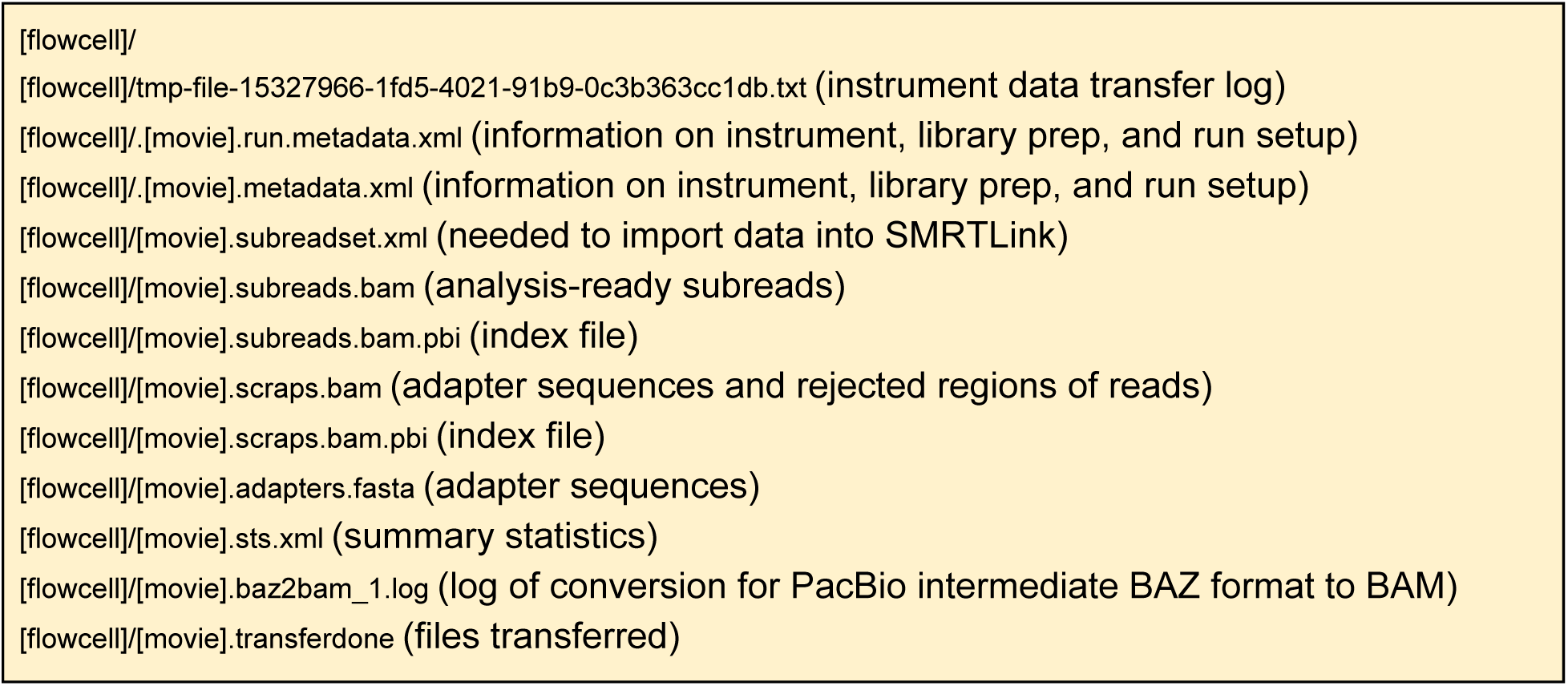
Raw data tar.gz directory structure.

~~~
samtools fasta [movie].subreads.bam | gzip -c
~~~

Reads were aligned to reference genomes GRCh37 with hs37d5 decoy^3^ and GRCh38 with hs38d1 decoy^4^. A representative subread per zero-mode waveguide (ZMW) was extracted with `pbsv` (pbsv version 2.0.0, https://github.com/PacificBiosciences/pbsv) and aligned to the reference with `minimap2` (version 2.11-r797, Li 2018). Per-movie alignments were merged into a single aligned BAM and indexed using `samtools` (version 1.3.1).

~~~
pbsv fasta [movie].subreads.bam | \
     minimap2 -t 8 -x map-pb -a --eqx -L -O 5,56 -E 4,1 -B 5 \
     --secondary=no -z 400,50 -r 2k -Y [reference].fa - | \
     samtools sort > [sample]_[movie]_[reference].bam
~~~

## Data Records

### Genomic Samples

The GIAB Han Chinese trio genomes are available as EBV-immortalized cell lines and DNA from Coriell (Table 1). Genomic DNA from the son is available as a NIST Reference Material (RM8393). RM8393 genomic DNA was prepared from a single homogeneous culture by Coriell specifically for the NIST reference material. The subjects are approved for “Public posting of personally identifying genetic information (PIGI)” by the Coriell and NIH/NIGMS IRBs. The study was approved by the NIST Human Subjects Protections Office and Coriell/NIGMS IRB.

### Sequence Data

The sequence data is available as raw data, sequences (fasta), and aligned reads (bam) at the NCBI GIAB ftp site (links below). The raw data are in the `raw_data` subdirectory as tar.gz files (Figure 1). The tar.gz files are named using the following naming convention [Cell UUID].tar.gz. The compressed data archives include subreads as bam files (BAM file format specifications http://samtools.github.io/hts-specs/SAMv1.pdf, PacBio bam file format specifications https://pacbiofileformats.readthedocs.io/en/5.1/BAM.html). Sequence data is available in the `PacBio_fasta` subdirectory as gzipped fasta files with the following naming convention `[movie].subreads.fasta.gz`. When base quality information is needed, e.g. read mapping, the subread bam files in the `raw_data` can be used. The aligned read data is located in the `PacBio_minimap2_bam` subdirectory. The aligned reads are provided as bam files along with their index.^5^ The bam file names use the following convention `[NIST ID]_PacBio_[REF ID].bam`, where `[REF ID]` indicates the reference genome that was used and is either GRCh37 or GRCh38. The raw data is archived in the NCBI Sequence Read Archive (SRA) under accessions NCBI SRA SRX4739017 for HG005 Biosample SAMN03283350, SRX4739121 for HG006 Biosample SAMN03283348, and SRX4739122 for HG007 Biosample SAMN03283349, under SRP047086 with BioProject PRJNA200694. A list of fasta files with the ftp paths for each sample can be obtained via a sequence index file.^6^

**Son** ftp://ftp-trace.ncbi.nlm.nih.gov/giab/ftp/data/ChineseTrio/HG005_NA24631_son/MtSinai_PacBio/

**Father** ftp://ftp-trace.ncbi.nlm.nih.gov/giab/ftp/data/ChineseTrio/HG006_NA24694-huCA017E_father/PacBio_MtSinai/

**Mother** ftp://ftp-trace.ncbi.nlm.nih.gov/giab/ftp/data/ChineseTrio/HG007_NA24695-hu38168_mother/PacBio_MtSinai/

## Technical Validation

The sequence dataset was characterized for number of reads, read length, coverage, mapping quality, and error rate. Mapped reads were used to characterize coverage, mapping quality, and error rate for the three samples. Metrics were calculated for reads mapped to GRCh37 using `minimap2` (see Methods for details) using `samtools stats`. Nearly three times the number of SMRT Cells were used in sequencing HG005 compared to HG006 and HG007 (Table 2) resulting in approximately twice the total number of reads (Table 2). Improved loading efficiency was observed when using the later v2.1 sequencing chemistry. The majority (39/46) of SMRT Cells from HG0005 were run with v2.0; whereas the majority (21/33) of SMRT Cells of the parental DNA was sequenced with v2.1. The polymerase did not change between v2.0 and v2.1 sequencing kits and therefore use of different sequencing kit is only expected to affect throughput and not error rates. Mean read length and N50 is similar across samples with mean subread lengths between 9.8 kb and 10.4 kb and N50 between 16.7 kb and 18.8 kb (Table 2, Figure 2). HG005 had approximately twice the coverage of HG006 and HG007 (Table 3, Figure 3). HG005 had ~15X coverage by reads >20kb and HG006 and HG007 had ~10X coverage (Figure 4). The mapping rate was higher for HG005 compared to the other two samples (88% vs 83%). For HG006 the MQ0 rate (MQ0 rate is the percent of the mapped reads with a mapping quality of 0) was higher than the other two samples (0.40% versus 0.36%, Table 3). The base pair error rate is around 15% for all three samples.

**Figure 2:**
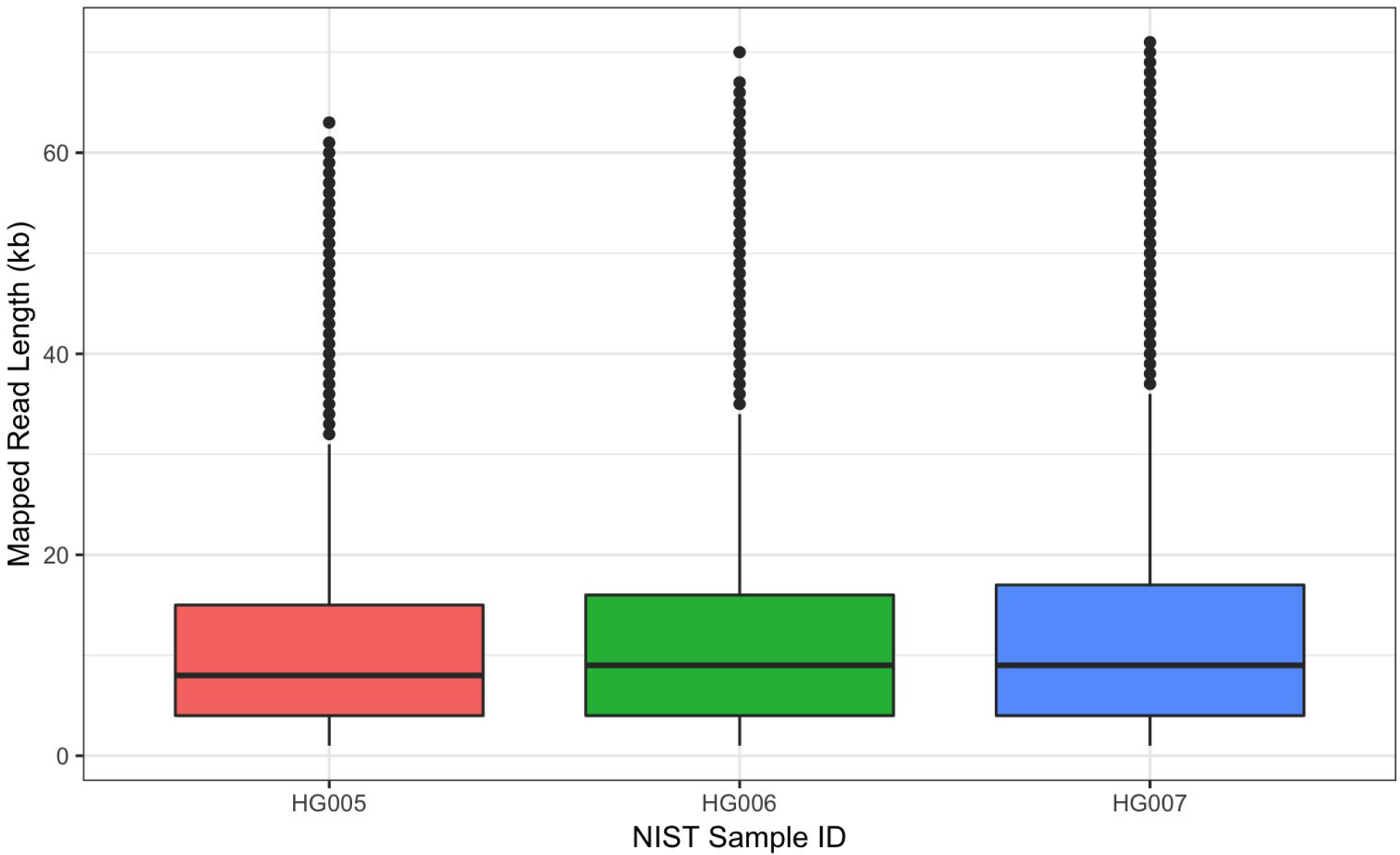
Read length distribution of mapped reads for the three samples. Boxplot horizontal bars indicate median read length, boxes the interquartile range, whiskers ± 1.5 ✕ interquartile range, and distribution outliers indicated with black points.

**Figure 3:**
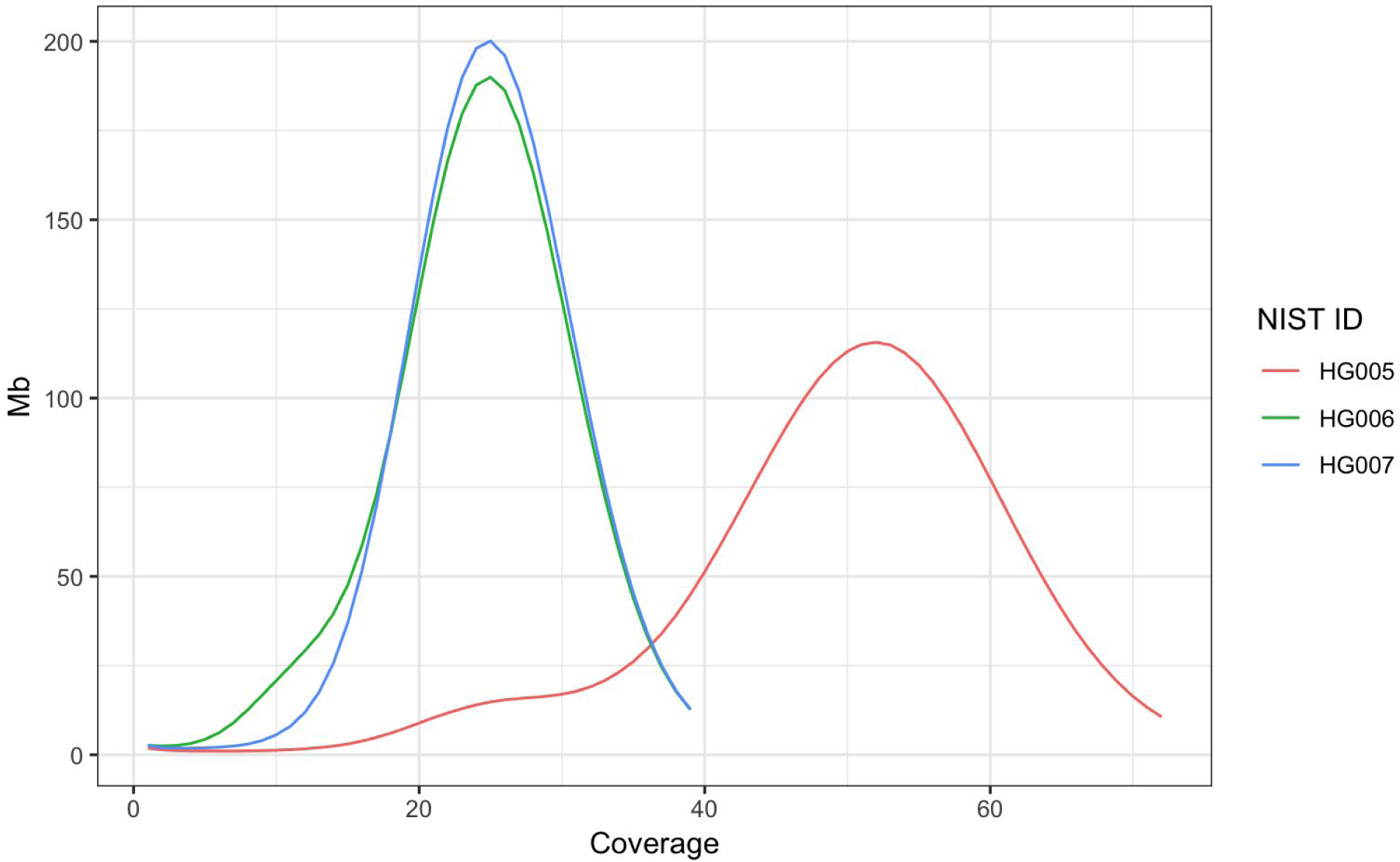
Distribution in the number of mapped Mb (10^6^ bases) by coverage.

**Figure 4:**
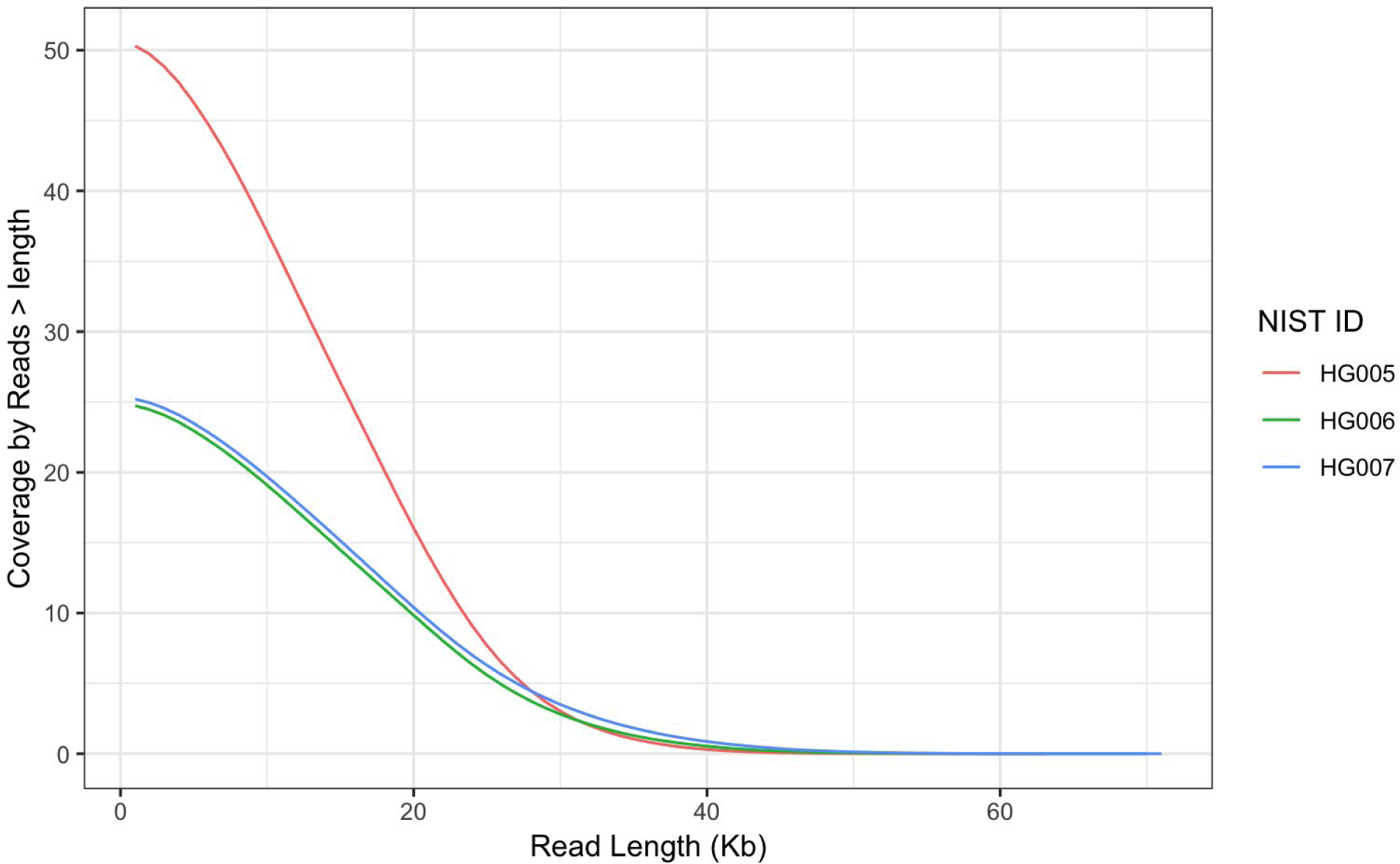
Distribution in the of coverage by mapped read length

**Table 2:**
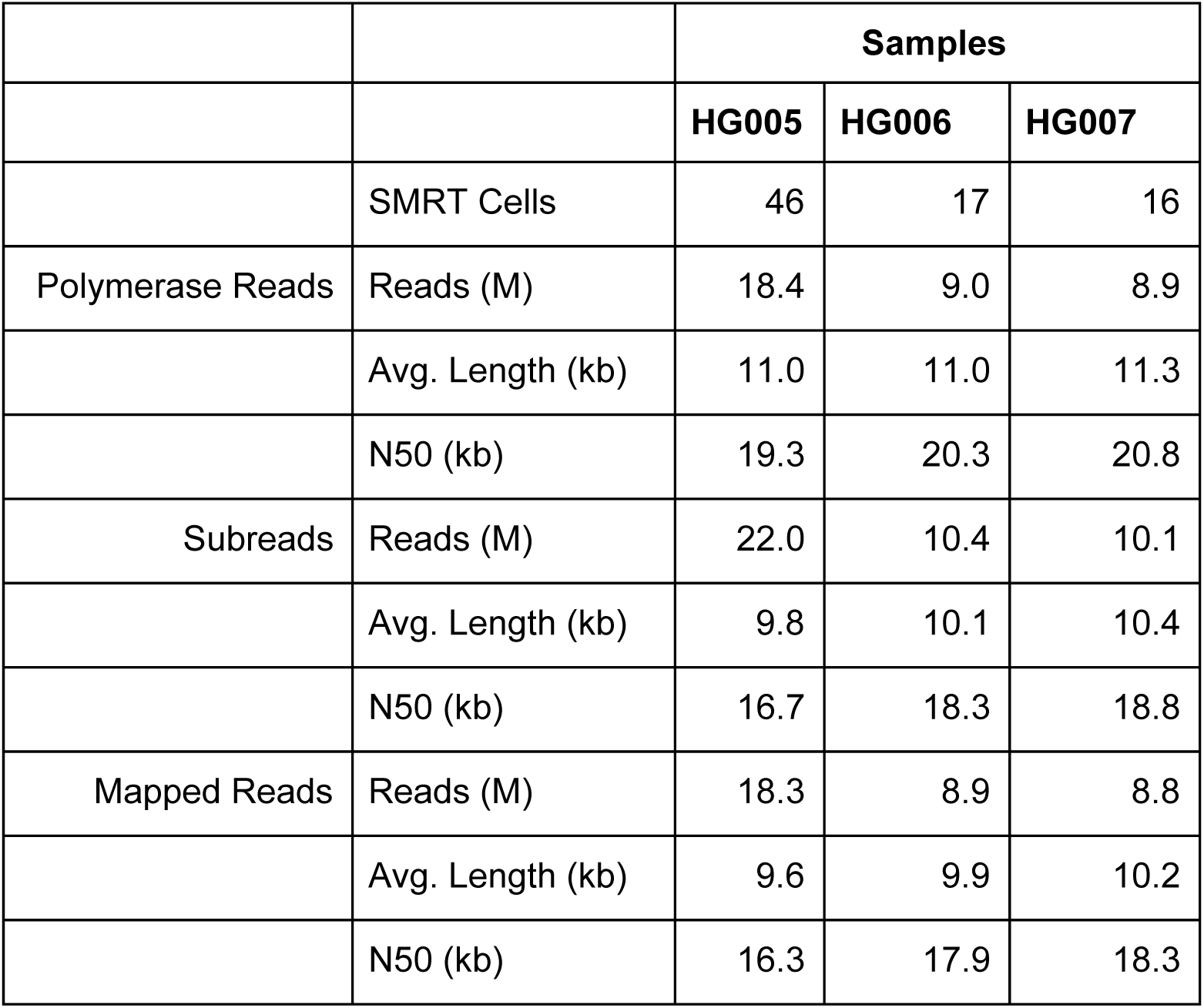
Pacific Biosciences Sequel run metrics. Read metrics provided for polymerase reads, subreads, and mapped reads. Subreads (inserts) are sequences between SMRTbell adapters, the polymerase reads include SMRTbell adapters, and mapped reads are subreads mapped to GRCh37. Length (kb) - mean read length. Half of the sequenced bases are in reads longer than the N50.

**Table 3:** Read mapping summary metrics. Read mapping metrics were calculated for reads mapped to GRCh37 using `minimap2`. Coverage is the mean number of reads mapped to each position in the genome. Mapping rate is the number of mapped reads/ total number of subreads. MQ0 rate is the percent of the mapped reads with a mapping quality of 0 (i.e., reads that map equally well to multiple genomic locations). The error rate is the number of mismatches and gaps (insertions and deletions) in the alignment divided by the number of mapped bases. The number of mapped bases was calculated from the cigar string. Metrics were calculated from bam files using the `samtools stats` command.

| Sample | Coverage | Mapping Rate | MQ0 Rate | Error Rate |
| --- | --- | --- | --- | --- |
| HG005 | 56.9 | 87.5% | 0.37% | 14.7% |
| HG006 | 28.5 | 82.9% | 0.40% | 14.9% |
| HG007 | 29.1 | 83.3% | 0.36% | 15.1% |

## Usage Notes

The data presented here can be used to evaluate different bioinformatic methods including small and structural variant calling, haplotyping, and genome assembly. All data from the Genome in a Bottle project are available without embargo, and the primary location for data access is ftp://ftp-trace.ncbi.nlm.nih.gov/giab/ftp. The data is also available as an Amazon Web Services Public Datasets repository with ‘s3://giab’ as bucket name and in the NCBI BioProject (http://www.ncbi.nlm.nih.gov/bioproject/200694). Additional information regarding GIAB project can be obtained from GIAB github site (https://github.com/genome-in-a-bottle/). Genome in a Bottle Consortium Analysis Team was formed to coordinate analyses. Analysis performed by the team are available on the ftp site.^7^ Analysis subdirectories generally use the following naming convention, `[Dataset Name]_[Tool]_[Date (MMDDYYYY)]`. Benchmark callsets are available ftp://ftp-trace.ncbi.nlm.nih.gov/giab/ftp/release/ for use in evaluating small variant calling pipelines (Zook et al. 2018). For best practices in evaluating small variant calling methods see Krusche et al. (2018). Methods and benchmark callsets for structural variant detection are in active development (Truvari - https://github.com/spiralgenetics/truvari, SVanalyzer - https://svanalyzer.readthedocs.io/en/latest/). No reference assemblies are available for these genomes and methods for evaluating genome assemblies in an active area of research.

## Data Citations

Zook, J.M. et al. NCBI SRA SRX4739017 (2018).

Zook, J.M. et al. NCBI SRA SRX4739121 (2018).

Zook, J.M. et al. NCBI SRA SRX4739122 (2018).

## Acknowledgments

Certain commercial equipment, instruments, or materials are identified to specify adequately experimental conditions or reported results. Such identification does not imply recommendation or endorsement by the National Institute of Standards, nor does it imply that the equipment, instruments, or materials identified are necessarily the best available for the purpose.

## Author contributions

RS, MLS, JMZ - experimental design, data processing, manuscript preparation

GD - SMRTbell template preparation and sequencing

HS, YCW, AMW - data processing, bioinformatics

NDO - data analysis, manuscript preparation

JT - data submission

CX - data submission and management, manuscript preparation

SS - project and data management

MS - project design and execution

All authors have reviewed and approved the manuscript.

## Footnotes

1 https://www.thejournalofprecisionmedicine.com/archive-manager/conference-report/

2 ftp://ftp-trace.ncbi.nlm.nih.gov/giab/ftp/data/ChineseTrio/

3 ftp://ftp.1000genomes.ebi.ac.uk/vol1/ftp/technical/reference/phase2_reference_assembly_sequence/hs37d5.fa.gz

4 ftp://ftp.ncbi.nlm.nih.gov/genomes/all/GCA/000/001/405/GCA_000001405.15_GRCh38/seqs_for_alignment_pipelines.ucsc_ids/GC_A_000001405.15_GRCh38_no_alt_plus_hs38d1_analysis_set.fna.gz

5 https://samtools.github.io/hts-specs/

6 https://github.com/genome-in-a-bottle/giab_data_indexes/blob/master/ChineseTrio/sequence.index.ChineseTrio_NIST_MtSinai_Pac_Bio_Sequel_fasta_09282018

7 ftp://ftp-trace.ncbi.nlm.nih.gov/giab/ftp/data/ChineseTrio/analysis/

